# DNA-PKcs kinase activity stabilizes the transcription factor Egr1 in activated immune cells

**DOI:** 10.1101/2021.06.04.446996

**Authors:** Zachary J Waldrip, Lyle Burdine, David K Harrison, Ana Clara Azevedo-Pouly, Aaron J. Storey, Olivia G. Moffett, Samuel G. Mackintosh, Marie Schluterman Burdine

## Abstract

DNA-dependent protein kinase catalytic subunit (DNA-PKcs) is known primarily for its function in DNA double-stranded break repair and non-homologous end joining (NHEJ). However, DNA-PKcs also has a critical yet undefined role in immunity impacting both myeloid and lymphoid cell lineages spurring interest in targeting DNA-PKcs for therapeutic strategies in immune-related diseases. To gain insight into the function of DNA-PKcs within immune cells, we performed a quantitative phosphoproteomic screen in T cells to identify first order phosphorylation targets of DNA-PKcs. Results indicate that DNA-PKcs phosphorylates the transcription factor Egr1 (early growth response protein 1) at S301. Expression of Egr1 is induced early upon T cell activation and dictates T cell response by modulating expression of cytokines and key costimulatory molecules. Mutation of serine 301 to alanine via CRISPR-Cas9 resulted in increased proteasomal degradation of Egr1 and a decrease in Egr1-dependent transcription of IL2 (interleukin-2) in activated T cells. Our findings identify DNA-PKcs as a critical intermediary link between T cell activation and T cell fate and a novel phosphosite involved in regulating Egr1 activity.

## Introduction

The canonical function for DNA-dependent protein kinase catalytic subunit or DNA-PKcs is in the sensing and repair of DNA double-strand breaks (DSB) through non-homologous end joining (NHEJ). However, all vertebrates harboring kinase loss of function mutations in DNA-PKcs present with a severe immunodeficient phenotype with defects in antibody production and impaired B and T cell maturation.^1–4^ These defects have been primarily attributed to the function of DNA-PKcs in V(D)J recombination which is required for antibody and receptor diversity in adaptive immune cells.^5^ Interestingly, this enzyme is robustly expressed in mature lymphocytes and consistently activated by various lymphocyte stimulants.^6,7^ This emphasizes a function for DNA-PKcs in the mature immune system that is yet to be clearly defined.

T cells are a key component of the adaptive immune response providing long term protection against evading pathogens. Uncontrolled or defective T cell activity, however, can have deleterious effects including transplant graft rejection, graft versus host disease and a plethora of autoimmune diseases.^8,9^ Therefore, understanding molecular mechanisms that regulate T cell activity are critical for the development of novel therapeutics to prevent/treat T cell-mediated disorders. T cell receptor (TCR) activation induces signaling cascades that regulate T cell proliferation, survival and differentiation. The end result is widely dependent on the activation of transcription factors that promote expression of cytokines and chemokines that, depending on the level and combination, can have varying effects on T cell response. For instance, graded expression of the T-box transcription factor T-bet in naïve CD4+ T cells coordinates helper (Th) 1 or T follicular helper (Tfh) cell differentiation with higher levels driving a Th1 cell fate.^10^ It is becoming clear that DNA-PKcs strongly influences T cell activity, as well as other immune cells, through regulation of transcription factor expression. In CD4+ T cells, following TCR activation, DNA-PKcs regulates expression of both T-bet and Gata3 highlighting it as a master regulator of Th1 and Th2 differentiation.^11,12^ Our laboratory recently reported that DNA-PKcs also controls expression of the p65 subunit of NF-κB in activated T cells and loss of DNA-PKcs activity significantly reduces expression of NF-κB target genes including Interleukin (IL)-6.^13^ Ferguson et al. determined that following viral DNA detection, DNA-PKcs drives activation of the innate immune response by directly binding the transcription factor interferon (IFN) regulator factor-3 (Irf-3) and promoting its translocation into the nucleus to induce cytokine gene expression.^14^ Similarly, our studies indicate that DNA-PKcs plays a pivotal role in the calcineurin-mediated translocation of NFAT to the nucleus. Inhibition of DNA-PKcs blocked calcineurin activity thereby preventing the translocation of NFAT to the nucleus and expression of cytokine IL2.^15^ Herein, we report that DNA-PKcs also regulates expression of the immediate early response gene (IEG) Egr1 (early growth response 1), a transcription factor critical for cytokine production.^16–18^ IEG genes such as Egr1 are transcribed within minutes of TCR stimulation to rapidly turn on transcription of genes needed for immune cell function.^19,20^ This includes genes like *NFKB, ELK*, and *NFAT* which can be activated quickly through degradation of inhibitors or through post-translational modifications via MAP kinase cascades.^21–23^ IEGs, therefore, are responsible for coaxing T cells down specific response pathways predetermined by the type of immunogenic stimulus encountered. We identified Egr1 to be a phosphorylation target of DNA-PKcs. Inhibition of DNA-PKcs or mutation of serine 301 of Egr1 resulted in significant downregulation of Egr1 protein leading to reduced secretion of IL2. Regulation of early signaling effectors like Egr1 suggests that DNA-PKcs functions as a critical link between TCR stimulation and subsequent gene transcription capable of guiding T cell signaling towards specific outcomes.

## Results

### Egr1 is phosphorylated by DNA-PKcs following stimulation of T cells

Given the diversity of signaling events in which DNA-PKcs is involved, we sought to identify potential DNA-PKcs phosphorylation targets involved in T cell activation. To accomplish this goal, we performed a quantitative proteomic mass spectrometry screen for phosphoproteins utilizing TMT (tandem mass tag, Thermo) technology. This was accomplished using human T cells (Jurkat) stimulated with phorbol myristate acetate and phytohemagglutinin (PMA and PHA, respectively) and treated with or without NU7441, a highly specific small molecule kinase inhibitor of DNA-PKcs.^24,25^ A phospho-TMT analysis combines affinity enrichment of phosphorylated proteins/peptides with highly quantitative TMT isobaric reagents; thus, allowing for a highly quantitative and extensive analysis of phosphorylation events in a cell. We analyzed for differentially phosphorylated proteins between stimulated cells and stimulated cells pre-treated with NU7441. **Figure 1** contains a list of all phosphoproteins with a fold change of >10. We identified phosphorylation of DNA-PKcs to be downregulated providing validity to our screen given that DNA-PKcs is known to autosphorylate itself at numerous sites. One of the more prominent phosphopeptides mapped to the IEG transcription factor Egr1. The phosphopeptide shown in **Figure 1** was approximately 30-fold more prevalent in the non-inhibitor treated sample. Analysis of this phosphorylation site revealed that it falls within the DNA-PKcs kinase recognition motif, SQD/E.^26^ To further substantiate the potential biological significance of this site, a sequence alignment of this peptide in vertebrates revealed a very high degree of conservation from zebrafish to humans (**Table 1**). As far as we know, phosphorylation of this serine residue (S301) has not previously been described.

**Figure 1.**
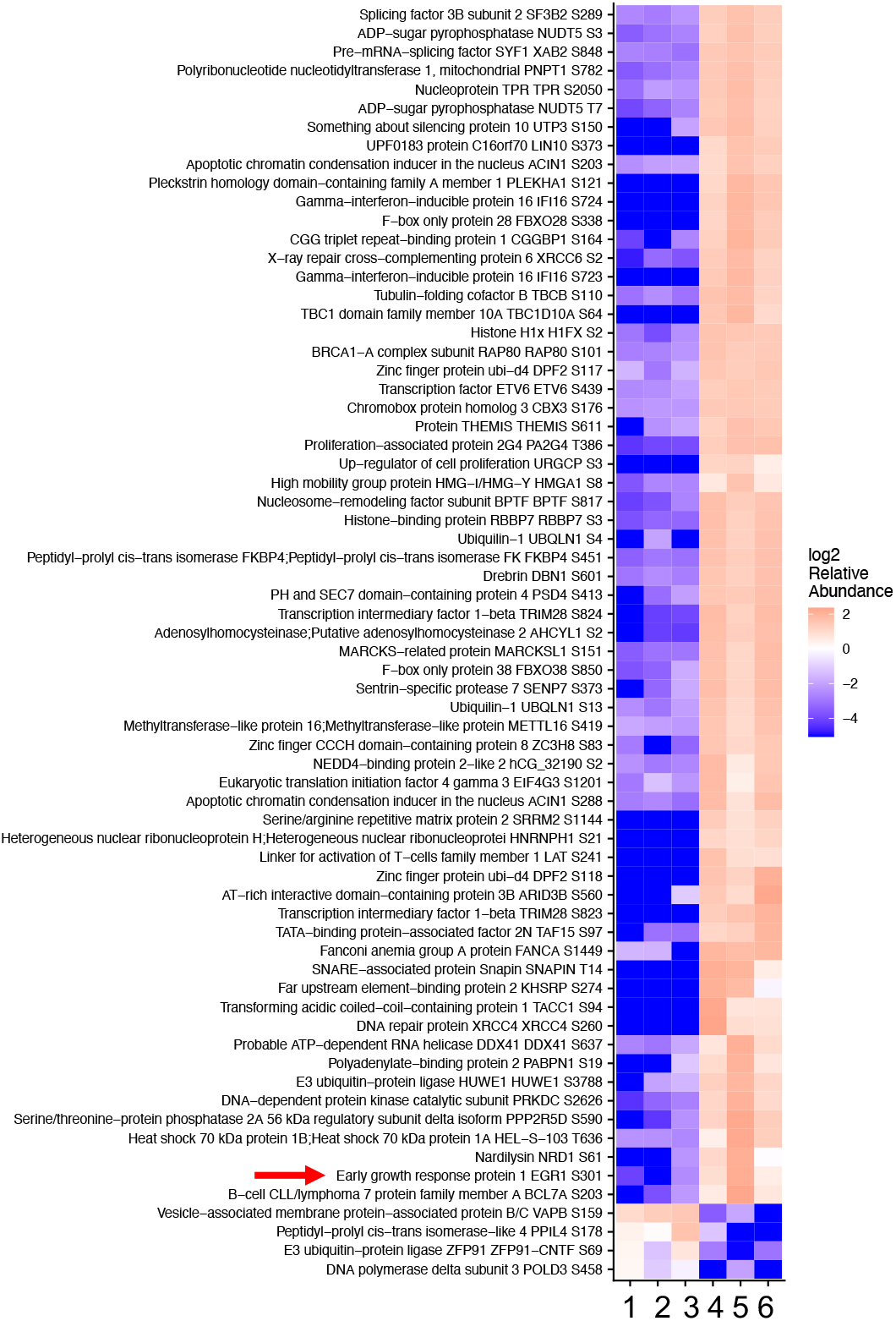
Heatmap of differentially regulated phospho-proteins in T cells treated with or without DNA-PKcs inhibitor. T cells were activated with PHA/PMA and treated with (samples 1-3) or without (samples 4-6) NU7441 for 6 hours and analyzed by mass spectrometry for comparison of differential abundance across groups. Heatmap contains all proteins with a fold change >10.

**Table 1.**
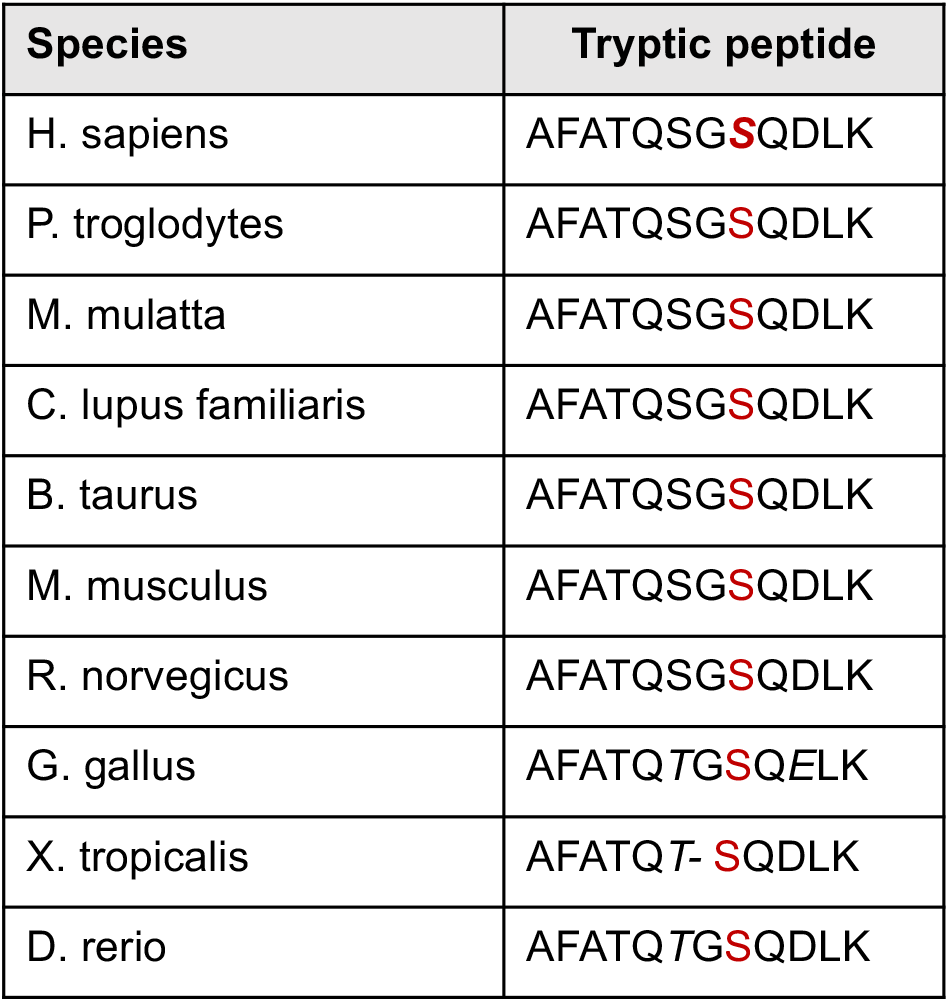
Egr1 S301 phosphorylation site detected by mass spectrometry is highly conserved in vertebrate animals. https://www.ncbi.nlm.nih.gov/homologene/56394

### Inhibition of DNA-PKcs kinase activity reduces Egr1 protein expression

Western blotting revealed that Egr1 was highly induced upon stimulation (**Figure 2**). However, Egr1 levels were markedly lower in NU7441-treated human T cells suggesting that DNA-PKcs kinase activity is required for Egr1 protein expression (**Figure 2A**). To confirm inhibition of DNA-PKcs activity by NU7441 in this assay, we probed for phospho-AKT at S473, a target of DNA-PKcs.^25^ Phosphorylation of S473 was significantly lower in NU7441 treated samples. To validate this result, we analyzed Egr1 expression patterns in total mouse splenocytes from wild type (WT) or *PRKDC* (gene for DNA-PKcs) knockout mice (KO) treated with NU7441 (**Figure 2B**) and in the embryonic kidney cell line HEK293 treated with shRNA to specifically knock down DNA-PKcs expression (**Figure 2C**). Inhibition of DNA-PKcs kinase activity significantly reduced Egr1 protein expression in both additional cell lines, indicating the robustness of this finding and confirming that this mechanism of regulation occurs in other cell lines that induce Egr1 expression to rapidly respond to cellular stimuli.

**Figure 2.**
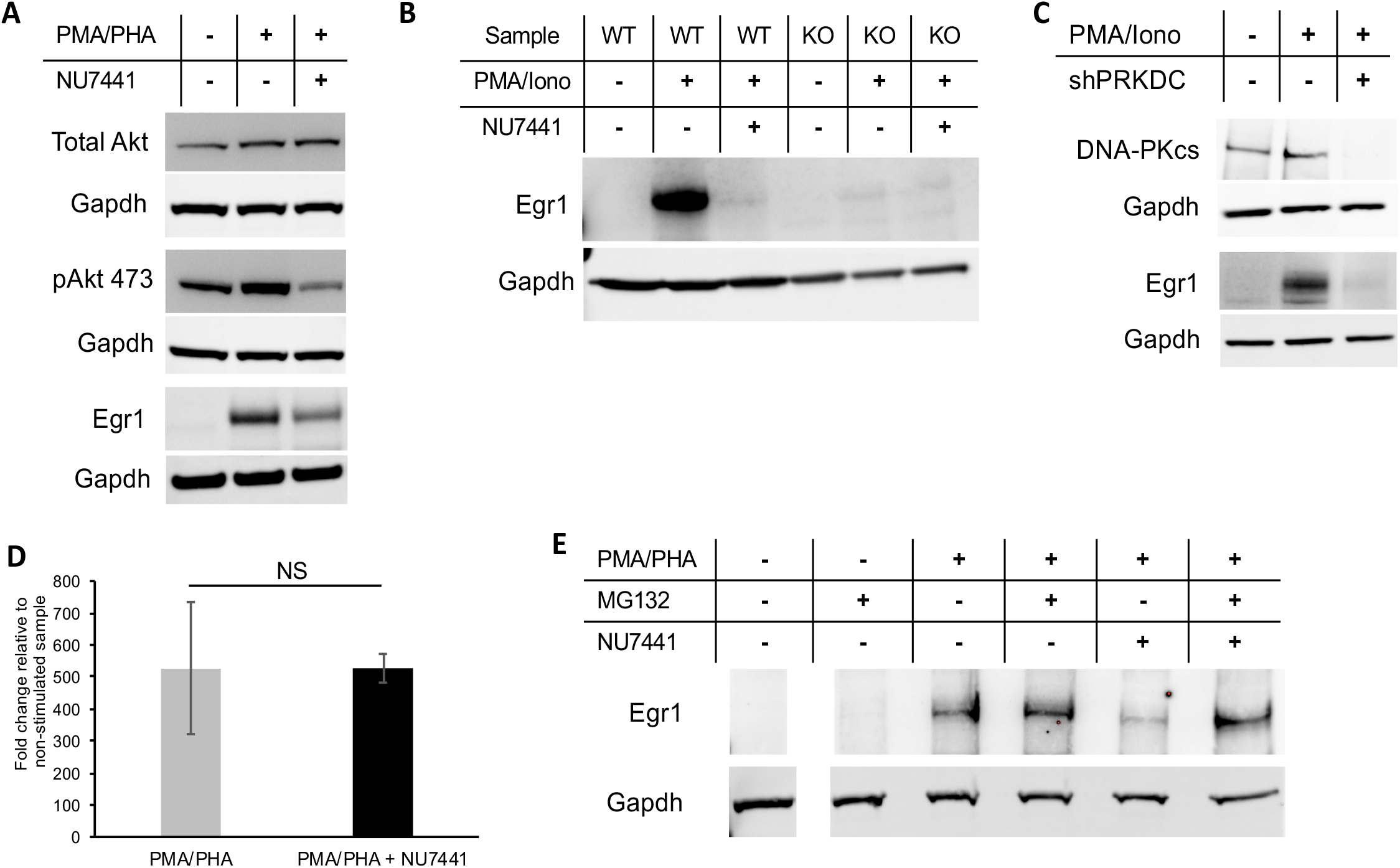
Protein and RNA expression patterns associated with Egr1 in multiple cell types. Egr1 expression pattern was analyzed by Western blotting in (A) Jurkat T cells, (B) total mouse splenocytes (WT indicates WT mouse and KO indicates PRKDC functional knockout mouse), and (C) HEK293 kidney cells which were chemically stimulated for 3 hours and treated with NU7441 as indicated. (D) Real-time qPCR analysis of EGR1 transcripts in Jurkat cells. Error bars represent standard error of the mean. NS indicates no significant difference. (E) Jurkat cells were treated as in (A) with the addition of the proteasome inhibitor MG132 and Egr1 was detected by Western blot.

*EGR1* is known to be tightly regulated at the transcriptional level and, like many other T cell-responsive transcription factors, the gene is highly induced upon T cell receptor or phorbol ester stimulation. To determine whether an effect of NU7441 on *EGR1* transcript levels explains the drop in Egr1 protein, qPCR was carried out in the presence or absence of NU7441. Transcript levels were unaffected (**Figure 2D**) by the loss of DNA-PKcs kinase activity. To assay whether the reduction in Egr1 protein was a result of proteasomal degradation, we analyzed Egr1 levels in NU7441-treated T cells in the presence of the proteasome inhibitor MG132. Our results indicate that proteasomal inhibition is able to restore the drop in Egr1 levels observed in the presence of NU7441 (**Figure 2E**).

### Phosphorylation of serine 301 of Egr1 is required for protein stability

To further analyze the functional relevance of S301 phosphorylation, plasmid-based FLAG-tagged Egr1 S301 mutants were expressed under the control of a constitutive promoter in HEK293 cells. An alanine mutant (S301A) as well as two phospho-mimetic mutants (S301D and S301E) were generated. Egr1 S301A protein levels were significantly decreased in contrast to both the WT protein (S301S) and the phospho-mimetic mutants S301D and S301E (**Figure 3A**). In fact, the phospho-mimetic mutations appeared to enhance protein stability. Next, we generated an *EGR1* knockout (*EGR1Δ*) as well as an endogenous S301A mutant in Jurkat T cells using CRISPR genome editing. This yielded a similar result where a single amino acid mutation to S301A to block phosphorylation resulted in a significantly reduced level of Egr1 protein despite normal transcription (**Figure 3B**). Transcript levels of the S301A mutant gene were similar to wild type *EGR1* levels (Supporting Figure 1).

**Figure 3.**
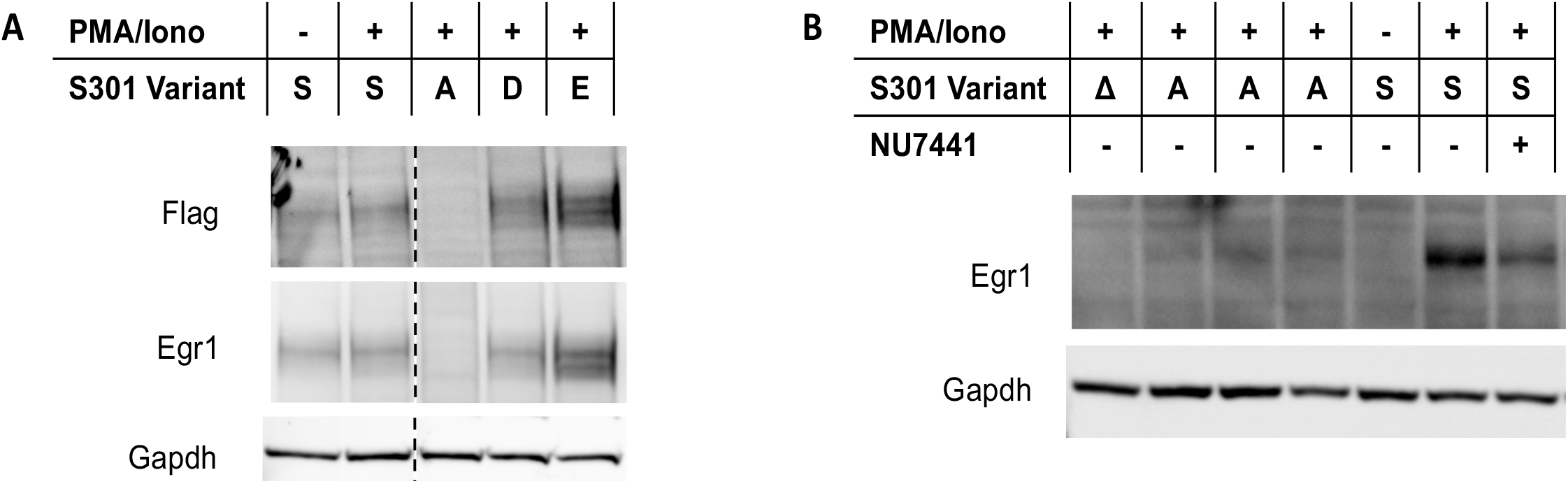
Effect of Egr1 phosphorylation on protein stability. (A) Four plasmid-based variants of Egr1-3xFlag at amino acid 301, as indicated by the single-letter amino acid abbreviation (A, D, E), were expressed in stimulated HEK293 cells. (B) Endogenous *EGR1* S301A and knockout mutants along with the wild type S301S strain were generated in Jurkat cells using CRISPR. Δ = *EGR1* CRISPR knockout, A = mutation generating S301A mutant (three separate clones are represented), S = WT *EGR1*.

### Loss of S301 phosphorylation abrogates production of IL2

Egr1 is a transcriptional regulator of the cytokine IL2. Therefore, to validate the relevance of this phosphorylation site to immune cell function, we analyzed expression of IL2. We used our CRISPR-generated *EGR1Δ* cell line to determine if the re-introduction of the Egr1 S301 variants (S301S, S301A, S301D, S301E) via electroporated plasmids would rescue IL2 expression. We hypothesized that the unstable S301A mutant would not fully restore or rescue IL2 levels. *EGR1Δ* cells were electroporated with one of five plasmids and stimulated 48 hours later using PMA and ionomycin for 6 hours. IL2 levels in the media were determined by ELISA. *EGR1Δ* samples transfected with plasmids expressing WT Egr1 S301S or the mutant S301D or S301E more than doubled their IL2 production relative to samples receiving control plasmid (**Figure 4**). In line with our findings, transfection of the S301A variant generated the lowest amount of IL2 rescue. IL2 levels from the cells expressing this variant were approximately 2.3-fold over the control, compared to 2.9 to 3.3 fold from cells expressing Egr1 S301S, S301D or S301E.

**Figure 4:**
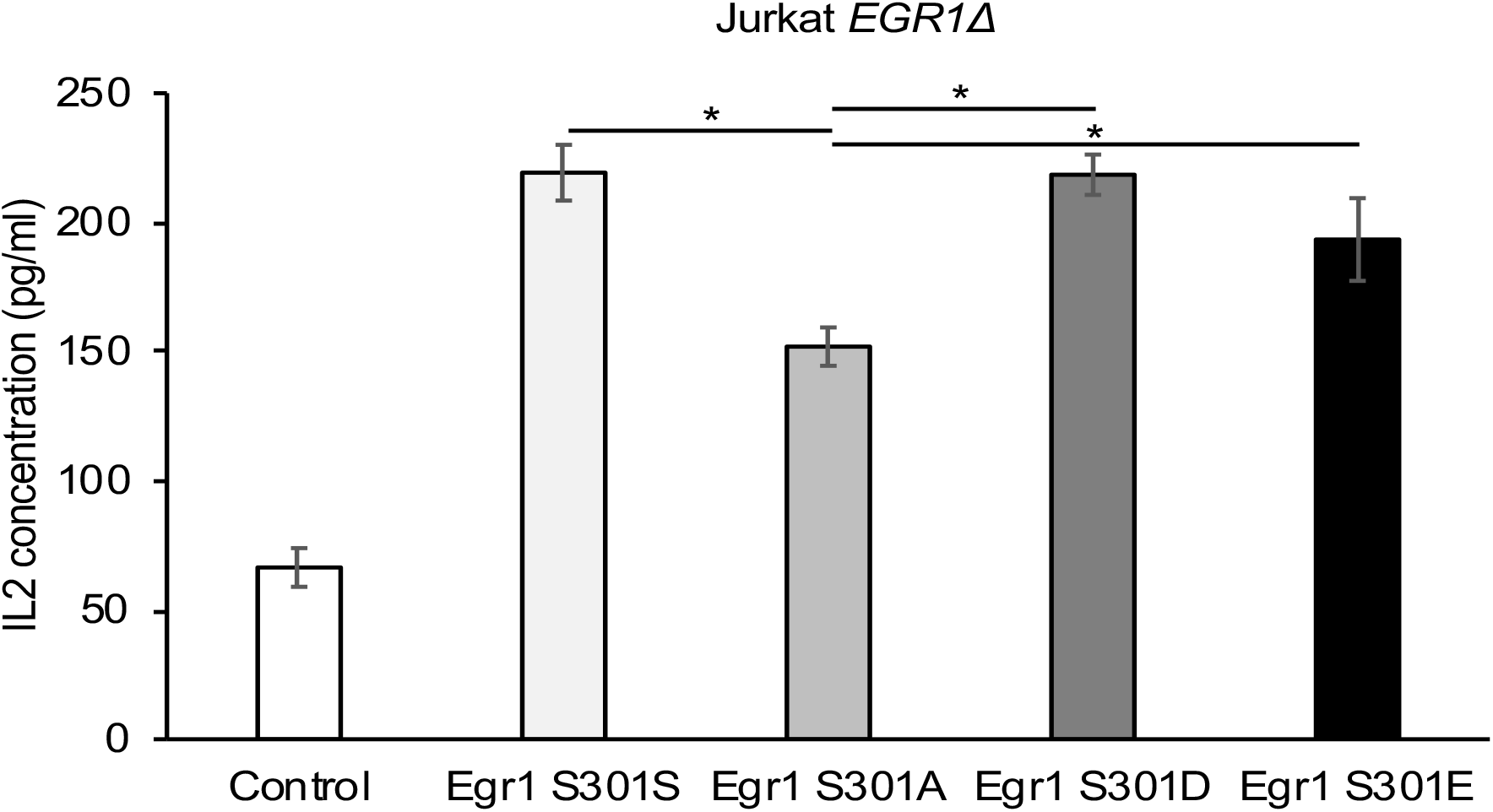
Effect of Egr1 phosphorylation on IL2 expression. IL2 concentrations were measured by ELISA in Jurkat *EGR1Δ* cells transfected by electroporation with plasmids expressing the indicated variant of Egr1. Control indicates transfection with a plasmid containing *GFP* in place of *EGR1*. Variability is represented by standard deviation of four replicates. * indicates p < 0.01 evaluated by T test.

## Discussion

While DNA-PKcs is a well-known mediator of double-stranded DNA damage through promotion of NHEJ, it is becoming increasing clear that it is also a critical regulator of the immune system. This is not a characteristic unique to DNA-PKcs. Other DDR kinases such as ATR and ATM have been linked to multiple processes in both the innate and adaptive responses.^27–30^ These functions are largely separate from their roles in NHEJ and HR (homologous recombination) which highlight a clear, yet largely undefined area of immune regulation. The goal of our study was to further understand mechanisms used by DNA-PKcs to govern T cell activation by uncovering novel target proteins. A mass spectrometry phosphoproteomic screen determined that the IEG transcription factor Egr1 is a robust phospho-target of DNA-PKcs. Phosphorylation of Egr1 at S301 by DNA-PKcs is required for protein stability and prevention of proteasomal degradation. IEGs are activated within thirty minutes of TCR activation and are decisive factors in mediating T cell responses to immunogenic stimuli.^19,31^ Identifying a role for DNA-PKcs in regulating expression and thus activity of Egr1 suggests that it has a much greater influence on immune response outcome than previously understood and could become a novel therapeutic target for a number of immune-related disorders. For instance, our previous study indicates that DNA-PKcs is a critical regulator of the T cell response to allogeneic antigens.^13^ Loss of DNA-PKcs activity prevented T cells from producing a host of inflammatory cytokines in response to alloantigen recognition resulting in reduced graft rejection *in vivo*. This suggests it may be a prime therapeutic target for the prevention of transplant rejection. While we show in those studies that the reduction in cytokine production from DNA-PKcs inhibition was partly due to a drop in protein expression of the NF*κ*B subunit p65, loss of Egr1 expression most likely was also involved in this outcome given that Egr1 promotes cytokine expression including IL2, a critical driver of transplant rejection.

We demonstrate that phosphorylation of Egr1 by DNA-PKcs prevents its degradation through the proteasomal pathway. Treatment with MG132 in the presence of NU7441 abrogated loss of Egr1 expression. Although the mechanisms involved in this effect are unknown and currently being investigated in our laboratory, this result has previously been observed for other DNA-PKcs targets. This includes estrogen receptor-*α*(ER-*α*) where interaction with DNA-PKcs resulted in phosphorylation at S118 which prevented proteasomal degradation.^32^ Our laboratory determined that DNA-PKcs controls NFAT-mediated transcription through indirect regulation of proteasomal degradation of the calcineurin inhibitor Cabin1.^6^ In contrast, DNA-PKcs has also been shown to promote ubiquitination of proteins for proteasome targeting. For instance, in order to arrest transcription at DSB sites, Caron et al. discovered that DNA-PKcs promotes ubiquitination of RNA polymerase II by HECT E3 ubiquitin ligase WWP2 thereby thwarting transcription.^33^ These studies clearly indicate that DNA-PKcs commonly uses the ubiquitin-proteasome pathway as a mechanism to control cellular functions. This includes the antigen-mediated T cell response where it uses this pathway to control expression of key transcription factors involved in T cell activation. Critical questions remain regarding mechanisms that induce DNA-PKcs activation following T cell stimulation and mechanisms used by DNA-PKcs to influence proteasome specificity. Identifying these mechanisms and further analyzing DNA-PKcs phosphorylation targets will uncover a novel area of immune regulation that further explains how T cells respond differently to varying stimuli and provide critical targets for novel therapy for immune-related diseases.

## Experimental Procedures

### Cell culture

Jurkat E6.1 cells and mouse splenocytes from BALB/c and NOD.CB17-Prkdc scid (#001303, #000651 respectively, Jackson Laboratory) were cultured in RPMI media supplemented with Penstrep antibiotics and 10% fetal bovine serum. HEK293 cells were cultured in EMEM media supplemented with Penstrep and 10% fetal bovine serum. Unless specified otherwise, cells were treated with NU7441 inhibitor (Selleckchem S2638) at a final concentration of 5 μM for 30 minutes prior to stimulation and harvested 3 hours post-stimulation. Stimulation was achieved with PMA (50 ng/ml) and PHA (1 μg/ml) or PMA (50 ng/ml) and Ionomycin (1 μg/ml).

### Western blotting

Samples for Western blots were processed by lysis in 0.5X RIPA buffer with protease inhibitors (Thermo Scientific #78425) and phosphatase inhibitors (Roche # 04906837001) followed by sonication in a QSonica Q800R3 with the settings 30% amplitude, 30s on/off, and 15 minute sonication time. Lysed samples were normalized by protein concentration determined using the bicinchoninic assay (BCA) (Thermo Scientific #23225). Samples were heated in LDS (lithium dodecyl sulfate) loading buffer then loaded into 4-12% bis-tris gels (Thermo Scientific #NW04122BOX). When blotting for DNA-PKcs, 3-12% tris-acetate gels were used (Thermo Scientific #EA0378BOX). Transfer to a PVDF membrane was completed using a Pierce Power Blotter system run at 25 V for 7 or 10 minutes (DNA-PKcs). Primary antibodies are as follows: Cell Signaling rabbit anti-Egr1 (44D5) #4154, Abcam mouse anti-DNA-PKcs [18-2] ab1832, Thermo Scientific mouse anti-Gapdh #MA5-15738, Cell Signaling rabbit anti-DYKDDDDK (FLAG) #2368, Cell Signaling rabbit anti-pAKT 473 #4060, and Cell Signaling rabbit anti-AKT #2938. Secondary antibodies are as follows: Thermo Scientific goat anti-mouse IgG (H+L) Alexa Fluor Plus 647 #A32728 and GE Healthcare donkey anti-rabbit HRP #NA934V. Imaging was done with a GE ImageQuant LAS4000.

### ELISA

Enzyme-linked immunosorbent assays were completed using and according to the instructions found with the ELISA MAX™ Deluxe Set Human IL-2 (BioLegend #431804).

### Mass spectrometry

Prior to analysis, two Jurkat cell samples were treated with PMA and PHA for 6 hours and one sample treated at 5 μM concentration with NU7441. Samples were harvested and lysed. Proteins were reduced, alkylated, and purified by chloroform/methanol extraction prior to digestion with sequencing grade trypsin and LysC (Promega). The resulting peptides were labeled using a tandem mass tag 10-plex isobaric label reagent set (Thermo) and enriched using High-Select TiO2 and Fe-NTA phosphopeptide enrichment kits (Thermo) following the manufacturer’s instructions. Both enriched and un-enriched labeled peptides were separated into 46 fractions on a 100 x 1.0 mm Acquity BEH C18 column (Waters) using an UltiMate 3000 UHPLC system (Thermo) with a 50 min gradient from 99:1 to 60:40 buffer A:B ratio under basic pH conditions, and then consolidated into 18 super-fractions. Each super-fraction was then further separated by reverse phase XSelect CSH C18 2.5 um resin (Waters) on an in-line 150 x 0.075 mm column using an UltiMate 3000 RSLCnano system (Thermo). Peptides were eluted using a 60 min gradient from 98:2 to 60:40 buffer A:B ratio. Eluted peptides were ionized by electrospray (2.2 kV) followed by mass spectrometric analysis on an Orbitrap Eclipse Tribrid mass spectrometer (Thermo) using multi-notch MS3 parameters. MS data were acquired using the FTMS analyzer in top-speed profile mode at a resolution of 120,000 over a range of 375 to 1500 m/z. Following CID activation with normalized collision energy of 31.0, MS/MS data were acquired using the ion trap analyzer in centroid mode and normal mass range. Using synchronous precursor selection, up to 10 MS/MS precursors were selected for HCD activation with normalized collision energy of 55.0, followed by acquisition of MS3 reporter ion data using the FTMS analyzer in profile mode at a resolution of 50,000 over a range of 100-500 m/z. Buffer A = 0.1% formic acid, 0.5% acetonitrile, Buffer B = 0.1% formic acid, 99.9% acetonitrile. Both buffers adjusted to pH 10 with ammonium hydroxide for offline separation. Protein identification, normalization and statistical analysis were performed as previously described by Storey et al.^34^

### Real-time PCR

RNA was purified with an Arum Total RNA mini kit (Bio-rad #732-6820). Reverse transcription was carried out using the iScript Advanced cDNA Synthesis kit (Bio-rad #1725037). Each PCR reaction was performed in technical duplicate and biological triplicate using SYBR green detection with a Bio-rad CFX96 Touch Real-time PCR detection system. Data was analyzed using the ΔΔCt method to determine relative concentrations of the EGR1 transcript normalized to TBP transcript levels. Primers for qPCR are as follows: EGR1 fwd – 5’ CAG CAC CTT CAA CCC TCA G, EGR1 rev – 5’ CAC AAG GTG TTG CCA CTG TT, TBP fwd – 5’ GCT GTT TAA CTT CGC TTC CG, TBP rev – 5’ CAG CAA CTT CCT CAA TTC CTT G

### Transfection

Transfection of HEK293 cells was accomplished using Lipofectamine 3000 (#L300000X) according to the manufacturer’s instructions. Transfection of Jurkat cells was accomplished by electroporation using the Lonza Nucleofector 4D system with the Amaxa SE Cell Line kit (#V4XC-1032). The manufacturer’s instructions were followed, and the pre-set Jurkat CL-120 program was used. In both cases, cells were grown for 48 hours post-transfection before use in experiments.

### Statistical analysis

Analysis of significance was done using standard t-test and expressed as the mean ± standard deviation. Assays were performed in triplicate. P ≤ 0.05 was considered significant.

## Funding

The project described was supported by the Center for Pediatric Translational Research NIH COBRE P20GM121293 and the Translational Research Institute (TRI) grant TL1 TR003109 through the National Center for Advancing Translational Sciences of the NIH. The content is solely the responsibility of the authors and does not necessarily represent the official views of the NIH.

## Acknowledgements

The mass spectrometry phosphoproteomics study was carried out in collaboration with the UAMS Discovery Proteomics Service Line under the guidance of Dr. Ricky Edmondson. Bioinformatics support was provided by director of the UAMS Bioinformatics Core Dr. Stephanie Byrum. We would also like to acknowledge the UAMS DNA Sequencing Core Facility at the Center for Microbial Pathogenesis and Host Inflammatory Responses for providing Sanger sequencing services.

---

**Supporting Figure 1:**
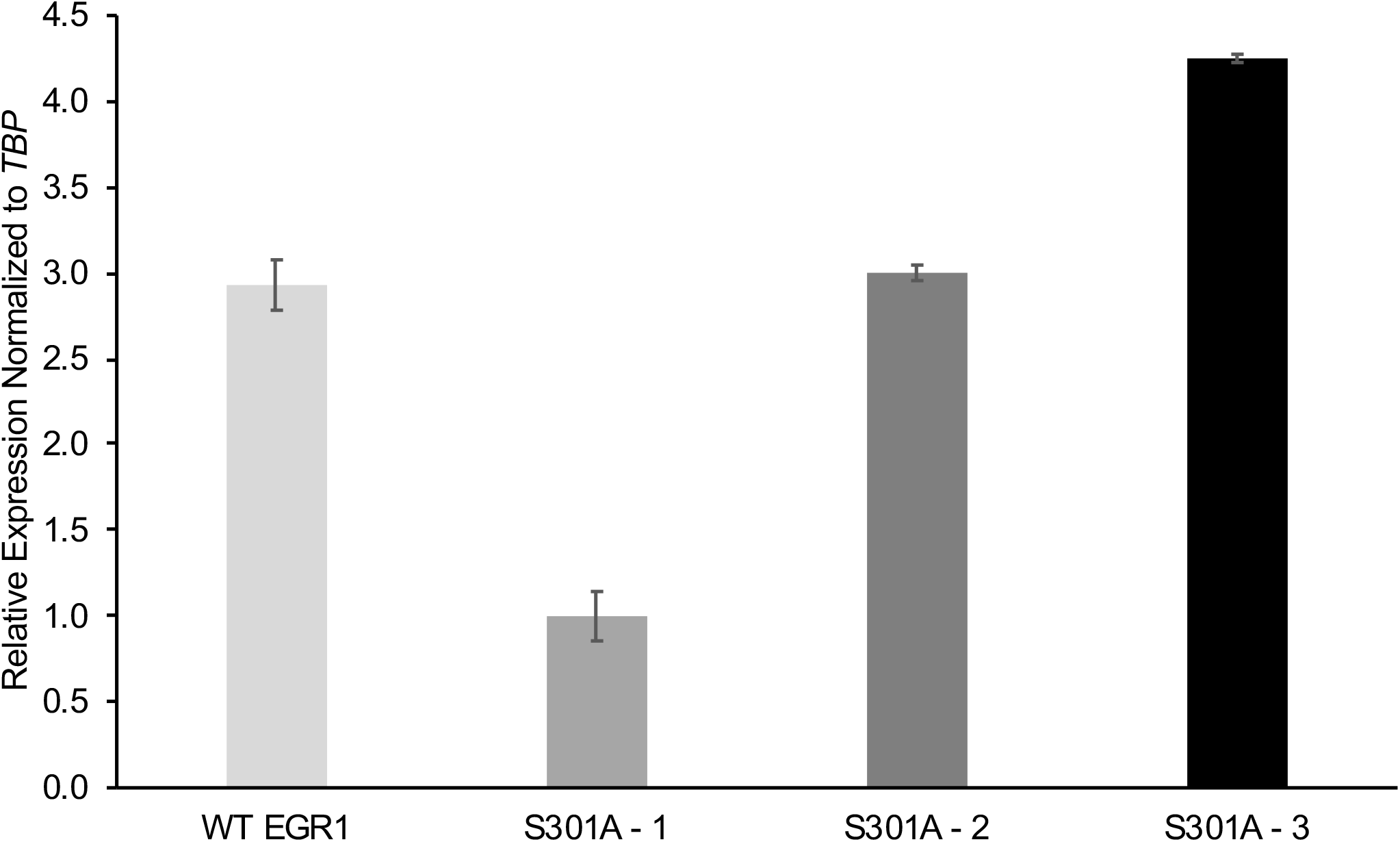
Egr1 S301 mRNA expression is similar to WT Egr1 mRNA expression. >Real-time qPCR analysis of *EGR1* transcripts from CRISPR-generated S301A mutants. Error bars represent standard error of the mean of technical replicates.

## Notes

### Competing Interest Statement

The authors have declared no competing interest.

## References

1. Davis AJ, Chen BPC, Chen DJ. DNA-PK: a dynamic enzyme in a versatile DSB repair pathway. DNA Repair (Amst). 2014;17:21–29. doi:10.1016/j.dnarep.2014.02.020

2. Pannunzio NR, Watanabe G, Lieber MR. Nonhomologous DNA end-joining for repair of DNA doublestrand breaks. J Biol Chem. 2018;293(27):10512–10523. doi:10.1074/jbc.TM117.000374

3. Taccioli GE, Amatucci AG, Beamish HJ, et al. Targeted disruption of the catalytic subunit of the DNA-PK gene in mice confers severe combined immunodeficiency and radiosensitivity. Immunity. 1998;9(3):355–366. http://www.ncbi.nlm.nih.gov/pubmed/9768755. Accessed January 8, 2018.

4. van der Burg M, van Dongen JJ, van Gent DC. DNA-PKcs deficiency in human: long predicted, finally found. Curr Opin Allergy Clin Immunol. 2009;9(6):503–509. doi:10.1097/ACI.0b013e3283327e41

5. Smider V, Chu G. The end-joining reaction in V(D)J recombination. Semin Immunol. 1997;9(3):189–197. doi:10.1006/SMIM.1997.0070

6. Kim Wiese A, Schluterman Burdine M, Turnage RH, Tackett AJ, Burdine LJ. DNA-PKcs controls calcineurin mediated IL-2 production in T lymphocytes. Trajkovic V, ed. PLoS One. 2017;12(7):e0181608. doi:10.1371/journal.pone.0181608

7. Ghonim MA, Pyakurel K, Ju J, et al. DNA-dependent protein kinase inhibition blocks asthma in mice and modulates human endothelial and CD4^+^ T-cell function without causing severe combined immunodeficiency. J Allergy Clin Immunol. 2015;135(2):425–440. doi:10.1016/j.jaci.2014.09.005

8. Dornmair K, Goebels N, Weltzien HU, Wekerle H, Hohlfeld R. T-cell-mediated autoimmunity: Novel techniques to characterize autoreactive T-cell receptors. Am J Pathol. 2003;163(4):1215–1226. doi:10.1016/S0002-9440(10)63481-5

9. Issa F, Schiopu A, Wood KJ. Role of T cells in graft rejection and transplantation tolerance. Expert Rev Clin Immunol. 2010;6(1):155–169. doi:10.1586/eci.09.64

10. Yu F, Sharma S, Edwards J, Feigenbaum L, Zhu J. Dynamic expression of transcription factors T-bet and GATA-3 by regulatory T cells maintains immunotolerance. Nat Immunol. 2015;16(2):197–206. doi:10.1038/ni.3053

11. Ghonim MA, Pyakurel K, Ju J, et al. DNA-dependent protein kinase inhibition blocks asthma in mice and modulates human endothelial and CD4+ T-cell function without causing severe combined immunodeficiency. J Allergy Clin Immunol. 2015;135(2):425–440. doi:10.1016/j.jaci.2014.09.005

12. Mishra A, Brown AL, Yao X, et al. Dendritic cells induce Th2-mediated airway inflammatory responses to house dust mite via DNA-dependent protein kinase. Nat Commun. 2015;6:6224. doi:10.1038/ncomms7224

13. Harrison DK, Waldrip ZJ, Burdine L, Shalin SC BM. DNA-PKcs Inhibition Extends Allogeneic Skin Graft Survival. Transplantation. 2020;Epub ahead(PMID:32890138).

14. Ferguson BJ, Mansur DS, Peters NE, Ren H, Smith GL. DNA-PK is a DNA sensor for IRF-3-dependent innate immunity. Elife. 2012;2012(1):47. doi:10.7554/eLife.00047

15. Wiese AK, Burdine MS, Turnage RH, Tackett AJ, Burdine LJ. DNA-PKcs controls calcineurin mediated IL-2 production in T lymphocytes. PLoS One. 2017;12(7). doi:10.1371/journal.pone.0181608

16. Shin HJ, Lee JB, Park SH, Chang J, Lee CW. T-bet expression is regulated by EGR1-mediated signaling in activated T cells. Clin Immunol. 2009;131(3):385–394. doi:10.1016/j.clim.2009.02.009

17. Li B, Power MR, Lin TJ. De novo synthesis of early growth response factor-1 is required for the full responsiveness of mast cells to produce TNF and IL-13 by IgE and antigen stimulation. Blood. 2006;107(7):2814–2820. doi:10.1182/blood-2005-09-3610

18. Pang Z, Raudonis R, McCormick C, Cheng Z. Early growth response 1 deficiency protects the host against pseudomonas aeruginosa lung infection. Infect Immun. 2020;88(1). doi:10.1128/IAI.00678-19

19. Mortlock S-A, Wei J, Williamson P. T-Cell Activation and Early Gene Response in Dogs. Mills K, ed. PLoS One. 2015;10(3):e0121169. doi:10.1371/journal.pone.0121169

20. Lee SM, Vasishtha M, Prywes R. Activation and repression of cellular immediate early genes by serum response factor cofactors. J Biol Chem. 2010;285(29):22036–22049. doi:10.1074/jbc.M110.108878

21. Oeckinghaus A, Ghosh S. The NF-kappaB family of transcription factors and its regulation. Cold Spring Harb Perspect Biol. 2009;1(4). doi:10.1101/cshperspect.a000034

22. Besnard A, Galan-Rodriguez B, Vanhoutte P, Caboche J. Elk-1 a Transcription Factor with Multiple Facets in the Brain. Front Neurosci. 2011;5(MAR):35. doi:10.3389/fnins.2011.00035

23. Lee JU, Kim LK, Choi JM. Revisiting the concept of targeting NFAT to control T cell immunity and autoimmune diseases. Front Immunol. 2018;9(NOV):2747. doi:10.3389/fimmu.2018.02747

24. Zhao Y, Thomas HD, Batey MA, et al. Preclinical Evaluation of a Potent Novel DNA-Dependent Protein Kinase Inhibitor NU7441. Cancer Res. 2006;66(10):5354–5362. doi:10.1158/0008-5472.CAN-05-4275

25. Dong J, Ren Y, Zhang T, et al. Inactivation of DNA-PK by knockdown DNA-PKcs or NU7441 impairs non-homologous end-joining of radiation-induced double strand break repair. Oncol Rep. 2018;39(3):912–920. doi:10.3892/or.2018.6217

26. Bednarski JJ, Sleckman BP. At the intersection of DNA damage and immune responses. Nat Rev Immunol. 2019;19(4):231–242. doi:10.1038/s41577-019-0135-6

27. Sun L-L, Yang R-Y, Li C-W, et al. Inhibition of ATR downregulates PD-L1 and sensitizes tumor cells to T cell-mediated killing. Am J Cancer Res. 2018;8(7):1307–1316. http://www.ncbi.nlm.nih.gov/pubmed/30094103. Accessed March 9, 2021.

28. Dillon MT, Bergerhoff KF, Pedersen M, et al. ATR inhibition potentiates the radiation-induced inflammatory tumor microenvironment. Clin Cancer Res. 2019;25(11):3392–3403. doi:10.1158/1078-0432.CCR-18-1821

29. Petersen AJ, Rimkus SA, Wassarman DA. ATM kinase inhibition in glial cells activates the innate immune response and causes neurodegeneration in Drosophila. Proc Natl Acad Sci U S A. 2012;109(11):E656–E664. doi:10.1073/pnas.1110470109

30. Riabinska A, Lehrmann D, Jachimowicz RD, et al. ATM activity in T cells is critical for immune surveillance of lymphoma in vivo. Leukemia. 2020;34(3):771–786. doi:10.1038/s41375-019-0618-2

31. Bahrami S, Drabløs F. Gene regulation in the immediate-early response process. Adv Biol Regul. 2016;62:37–49. doi:10.1016/j.jbior.2016.05.001

32. Medunjanin S, Weinert S, Schmeisser A, Mayer D, Braun-Dullaeus RC. Interaction of the double-strand break repair kinase DNA-PK and estrogen receptor-α. Mol Biol Cell. 2010;21(9):1620–1628. doi:10.1091/mbc.E09-08-0724

33. Caron P, Pankotai T, Wiegant WW, et al. WWP2 ubiquitylates RNA polymerase II for DNA-PK-dependent transcription arrest and repair at DNA breaks. Genes Dev. 2019;33(11-12):684–704. doi:10.1101/gad.321943.118

34. Storey AJ, Naceanceno KS, Lan RS, et al. ProteoViz: A tool for the analysis and interactive visualization of phosphoproteomics data. Mol Omi. 2020;16(4):316–326. doi:10.1039/c9mo00149b

